# The brain clock portal system: SCN-OVLT

**DOI:** 10.1101/2021.01.24.427962

**Authors:** Yifan Yao, Alana Taub, Joseph LeSauter, Rae Silver

## Abstract

Vascular portal systems are key structures, necessary for transporting products directly from the capillary bed of one region to the capillary bed of another region in high concentrations, without first returning to the heart. The only known portal systems in the brain is the hypophyseal-pituitary portal system, a structure necessary for survival and reproduction. Secretions from specific populations of hypothalamic neurons travel into fenestrated capillaries of the median eminence (ME) and thence drain into portal veins which break up into the secondary capillary plexus of the anterior pituitary. Neurons of the hypothalamic suprachiasmatic nucleus (SCN), locus of the brain’s master clock, also produces secretions deeply implicated in health and survival (Karatsoreos, 2019; Musiek & Holtzman, 2016). Here we describe a portal system connecting the SCN and organum vasculosum of the lamina terminalis (OVLT) - a circumventricular organ (CVO). CVOs lie around ventricles and lack a blood-brain barrier, enabling communication between the blood, brain, and cerebrospinal fluid. This “clock portal system” points to entirely new routes and targets for secreted signals, restructuring our understanding of brain communication pathways. Whether any of the remaining six CVOs in the mammalian brain bear portal systems is yet to be determined.

## Main

### Portal systems, diffusible signals and neurosecretion

The great majority of capillary beds drain into veins which then drain into the heart, rather than into another capillary bed. In mammalian brain, the hypothalamic-pituitary portal system, first demonstrated over 80 years ago by Wislocki and King (1936) is the only exception to this pattern. It took about a decade to establish the importance of the portal signaling pathway, achieved in Geoffrey Harris’s landmark monograph, *Neural Control of the Pituitary Gland (1948)*. About three decades after that, the existence of specific neurohormones that travel in this portal system led to Guillemin and Schally’s 1977 Nobel prize in physiology and medicine. The establishment of additional portal systems would launch a bonanza of new directions into research on the neurovasculature, a requirement for understanding connectomes in the brain (Kleinfeld et al., 2011).

Neurons of the circadian clock in the SCN, like other hypothalamic neurons, produce diffusible substances that impact behavior and the physiology. Several lines of evidence support the existence of SCN secretions. Transplants of SCN tissue rescue circadian rhythms of locomotor activity in arrhythmic SCN-lesioned host animals, with the period of the donor animal (Ralph, Foster, Davis, & Menaker, 1990), irrespective of the attachment site within the 3rd ventricle (3V) (Servière, Gendrot, LeSauter, & Silver, 1994) and are effective even when the grafted tissue is encapsulated in a polymer plastic that blocks fiber outgrowth (Silver, LeSauter, Tresco, & Lehman, 1996). Signals that diffuse from the SCN include transforming growth factor alpha (Kramer et al 2001), prokineticin2 (Cheng et al., 2002; Prosser et al., 2007) and cardiotrophin-like cytokine (Kraves & Weitz, 2006) as paracrine outputs and the peptides vasoactive intestinal polypeptide (VIP), arginine vasopressin (AVP) and gastrin releasing peptide (GRP) (Maywood, Chesham, O'Brien, & Hastings, 2011; Ono, Honma, & Honma, 2016). Despite substantial evidence of humoral signals, a challenge regarding their biological significance is how this very small nucleus could possibly produce sufficient product to orchestrate rhythms throughout the body, unless the target of those secretions lies very nearby (Schibler, Ripperger, & Brown, 2003). Here we identify a vascular pathway for communication of diffusible signals in a hypothalamic portal system connecting the SCN and a nearby CVO, the OVLT.

### Midline structures and vasculature

Because the vessels that course between the SCN and OVLT lie along the midline at the very base of the 3V, they are readily destroyed during physical sectioning and tissue processing for anatomical analyses. To retain the integrity of these midline structures, we prepared whole mounts of the brain, from the olfactory tubercle to the medulla using iDISCO (see Methods). Our light sheet microscopic scans captured the blood vessel packed in the brain midline, including the CVOs therein (OVLT and ME), and the AVP-stained SCN, lying between the CVOs (Fig.1a-c).

**Fig. 1.**
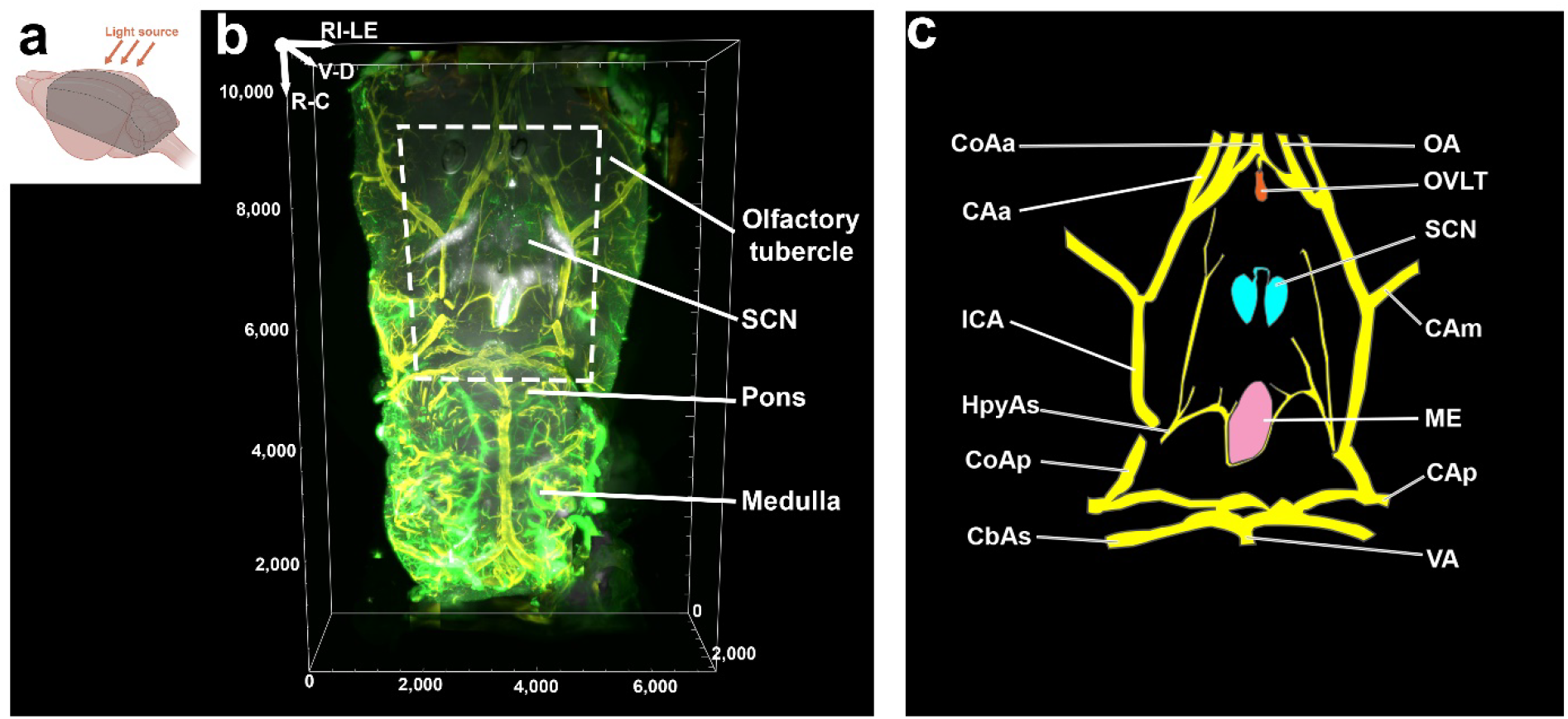
Ventral view of mouse brain from olfactory tubercle to medulla. **a,** Orientation of the scan. The gray area shows how the brain was blocked. The tissue was scanned horizontally with the light sheet parallel to the left-right axis (arrows). **b,** Ventral view of the brain from the olfactory tubercle to medulla in iDISCO cleared tissue labelled for AVP (white), collagen (green) and SMA (yellow). The scanned volume is from Bregma 2.10mm to −7.9mm in the rostro-caudal plane and ~3mm from the midline on each side. Voxel=5μm (R-C) x 5μm (LE-RI) x 5μm (D-V). l **c,** Line drawing shows structures within the dashed frame in b, for visualization of branches of the Circle of Willis (yellow), OVLT (orange), ME (pink) and SCN (blue). Abbreviations: CAa=anterior cerebral artery; CAm=medial cerebral artery; CAp=posterior cerebral artery; CbAs=superior cerebellar artery; CoAa=anterior communicating artery; CoAp=posterior communicating artery; HypAs=hypophyseal artery, superior branch; ICA=internal carotid artery; ME=median eminence; OA=olfactory artery; OVLT=vascular organ of lamina terminalis; SCN=suprachiasmatic nucleus; VA=vertebral artery. Fig. 1a was created with BioRender (https://biorender.com/). Reference axis denotes the orientation of the tissue and the same abbreviations were used for all images: R=rostral; C=caudal; V=ventral; D=dorsal; RI=right; LE=left. Scale unit=μm for all images unless specifically stated.

### SCN-OVLT capillary network

To examine the vasculature between the SCN and OVLT, we performed triple label immunochemistry using AVP to delineate the SCN, collagen as a general blood vessel marker, and smooth muscle actin (SMA) as an arterial marker and performed high power scanning. Each SCN has a pear-shaped main body. Right at the midline two prongs protrude rostrally from the medial most aspect of the SCN, and here lie blood vessels that travel between the SCN and the OVLT (Extended Data Fig. 1). The two prongs, termed SCN rostrum (SCNr), merge to form a bridge at the rostral-most pole of the nucleus (Extended Data Fig. 1 and Extended Data Fig. 3a, AVP panel). The capillary beds of the SCN and OVLT and the vessels lying between them were traced using Vesselucida360 (see methods). The results are shown in horizontal (Fig. 2bi-iii, and Fig. 2c) and coronal axes (Extended Data Fig. 2b). The traces point to several vessels emerging from the SCN rostrum (SCNr) to form a portal system running closely along the midline to the ventral OVLT. More laterally lying vessels are not part of this portal structure. The integrity of the tissue is confirmed by intact blood vessels and excellent correspondence with prior studies on the orientation and relative size of the hypothalamic region and ME, OVLT and SCN: rostrocaudally ~560μm for SCN main body and ~160μm for SCNr; dorsoventrally ~ 350μm; mediolaterally ~300μm; ~385μm between OVLT and SCN (Fig. 2a) which lie rostral to the ME and pituitary gland (Abrahamson & Moore, 2001; Allen Institute for Brain Science, 2004; La Perle & Dintzis, 2018; Lein et al., 2007; Varadarajan et al., 2018).

**Fig. 2.**
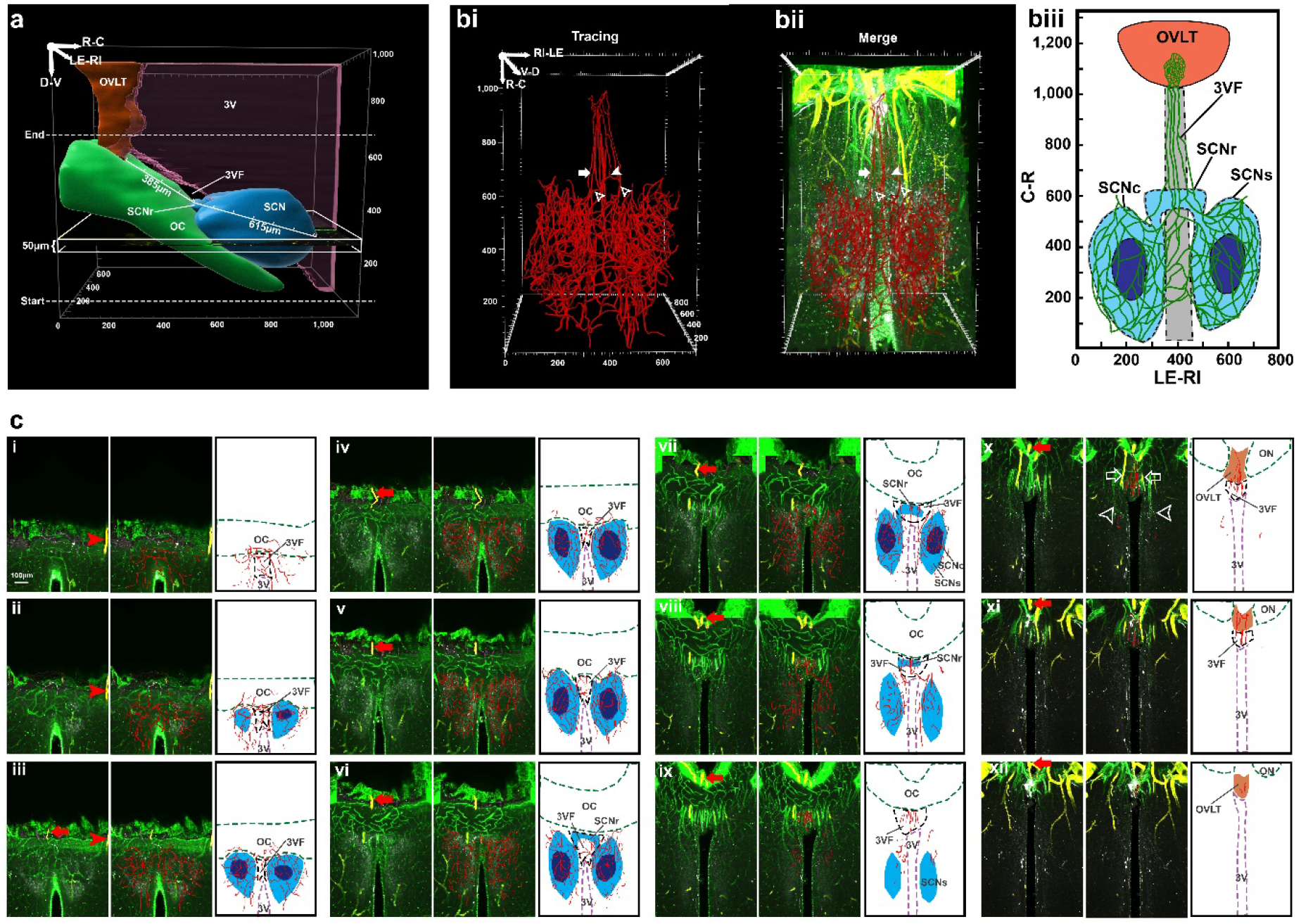
Capillaries vessels connecting SCN and OVLT in near-horizontal view. **a,** For analysis of the vasculature in the areas of interest, the OVLT (orange), SCN (blue), 3V (magenta) and OC (green) locations are shown. The white rectangle shows the orientation of the near-horizontal serial scans made through these regions, starting from the base of the brain (shown in c). **bi-iii,** Tracings of blood vessels in the SCN and OVLT region. **bi,** Tracings show capillary connection between the SCN and OVLT. **bii,** Tracings were merged with the image of the immunochemically stained collagen and SMA. In bi and ii, solid arrow=capillaries join midline blood vessel bundles; solid arrowhead=anastomoses; open arrowheads=capillaries of the SCN. **biii,** Schema of capillary vessels connection between OVLT and SCN in horizontal view. **c,** Serial optic slices of vasculature between SCN and OVLT. The plates show triplets of images as follows: left panel=merged AVP (white), collagen (green) and SMA (yellow); red arrows and arrowheads are place markers indicating landmarks for orientation in adjacent slices. Middle panel=blood vessel traces are superimposed on immunochemical results of the left panel; Right panel=drawing identifying structures in the middle panel. OVLT=orange; SCNs and SCNr=light blue, SCNc=dark blue. Details of the serial plates are as follows: **ci-v,** The left and right SCN capillary beds form anastomoses via capillaries traveling beneath the 3V. **cvi-viii**, The traced blood vessels near the SCNr form anastomoses. **cix-xi,** The portal blood vessels (red traces, middle panel) lie at the midline. **cx,** Veins that are not part of the portal system (middle column, white arrowheads) and arteries (white arrows) lie more laterally. **cxi-xii,** The midline blood vessels enter OVLT from its ventral side. Slice thickness=50μm. Abbreviations: 3VF=floor of the third ventricle; SCNc=SCN core; SCNr=SCN rostrum; SCNs=SCN shell; remaining abbreviations as in Fig. 1.

For a fine-grained analysis of the midline blood vessels, we made serial optical slices in (near-) horizontal (Fig. 2c) and in coronal orientations (Extended Data Fig. 2b). The main body of the two SCN nuclei are connected by capillaries that lie in the 3rd ventricle floor (3VF) above the optic chiasm (OC) thus sharing a common blood supply (Horizontal, Fig. 2ci-v; Coronal, Extended Data Fig. 2bi-iv). Additionally, capillaries form anastomoses along the SCNr (Horizontal, Fig. 2bi-iii, 2cvi-viii; Coronal, Extended Data Fig. 2bv).

Most importantly, the portal system between the SCN and OVLT is seen clearly at the rostral-most tip of the nucleus, where the left and right prongs of the SCN connect to form a bridge. The portal capillary veins lie right near the midline, in the glia limitans of 3VF above the optic chiasm (Horizontal, Fig. 2cix-xi; Coronal, Extended Data Fig. 2bvi-vii) and reach the OVLT at its ventral-most aspect (Horizontal, Fig. 2xi-xii; Coronal, Extended Data Fig. 2bviii).

### Comparison of SCN shell and core vasculature

The core and shell of the SCN have different functions in orchestrating rhythmicity. The core receives retinal input from the retinohypothalamic tract, and is important in synchronizing the clock to the local environment. Core neurons project to the shell, the locus of AVP-expressing cells. AVP is a major output and synchronization signal of the SCN (Colwell, 2011; Shan et al., 2020). Images of optical slices of cleared tissue (Fig. 3a) and confocal microscopic images of immunochemically stained material (Extended Data Fig. 3a) in three orientations through the mid-SCN suggest a denser and more complex vascular network in the SCN shell (SCNs) than in the SCN core (SCNc). This is confirmed quantitatively in tracings of collagen labeled vasculature (Fig. 3bi, ii; Extended Data Fig. 3c) and by the density of branching nodes in the SCNc and SCNs (Fig. 3biii). The greater complex capillary network in the SCNs is consistent with its greater role in vascular signaling.

**Fig. 3.**
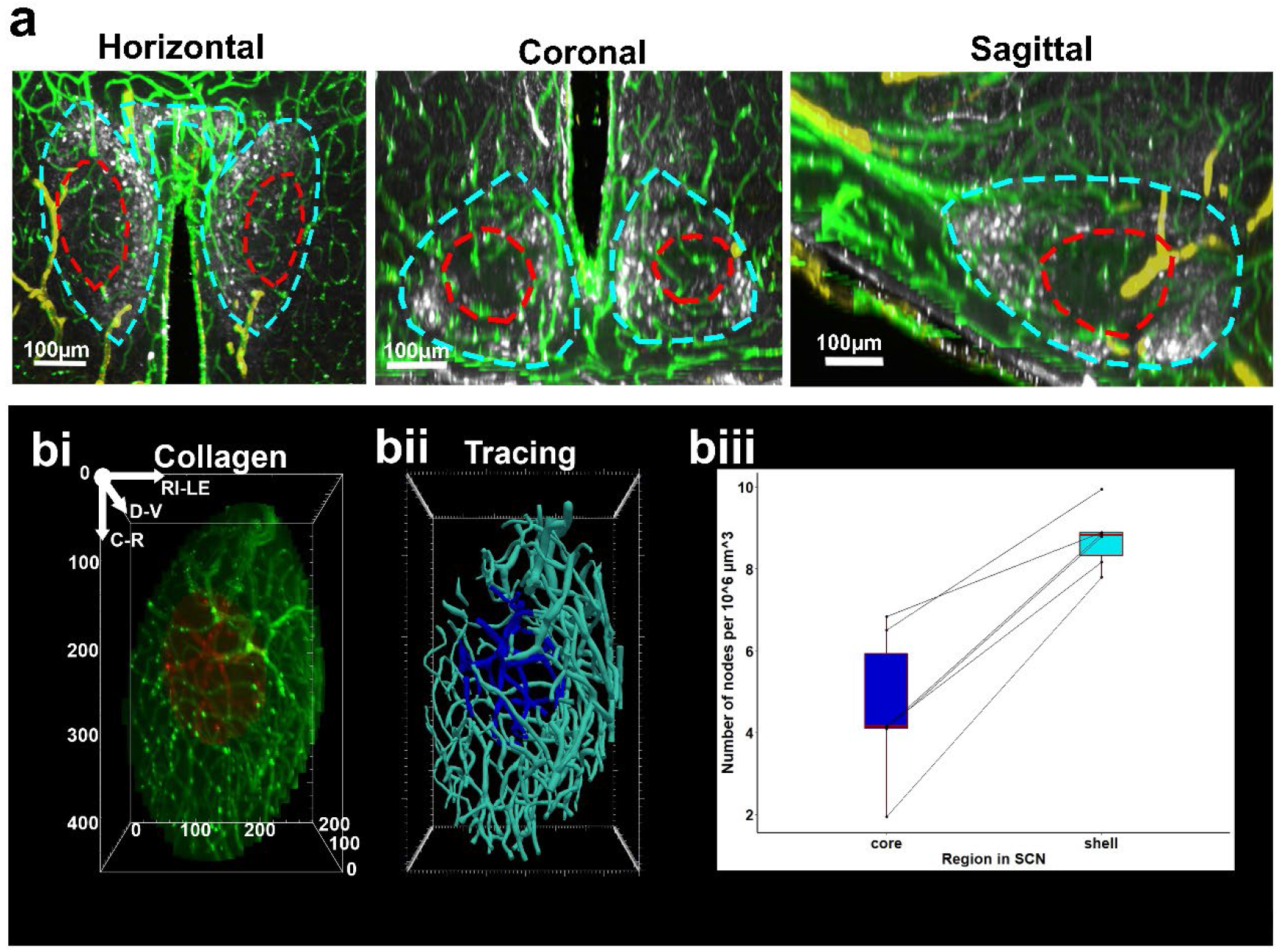
Capillary network of SCN core vs. shell in a cleared brain. **a,** Optical slices of a representative SCN in horizontal, coronal and sagittal orientation suggest a denser and more complex vasculature in the AVP-abundant shell (dashed blue outline) vs. core (dashed red outline). Slice thickness=50μm. **bi-iii,** Vasculature network analysis for the sample shown in Fig 2a. **bi,** Collagen staining of core (red) and shell (green) blood vessels **bii,** Tracings of SCN core (dark blue) and shell (light blue) blood vessels. **biii,** SCN core and shell branching density. SCN shell, 8.74±0.74 per 10^6^ μm^3^ vs. SCN core 4.61±1.81 per 10^6^ μm^3^; two-tailed paired t-test, t(5)=7.81, *p*=0.00055.

### Hypothalamic portal systems

Schematics summarizing the major features of the SCN-OVLT and the hypophyseal-pituitary portal systems are shown in Fig. 2biii and Fig. 4. The horizontal view (Fig. 2biii) emphasizes the capillary plexuses of the core and shell SCN, and the portal vessels that course between the SCN and OVLT. These vessels lie along the 3VF and are readily torn when the tissue is physically sectioned for anatomical analyses, but have survived intact in our cleared material.

**Fig. 4.**
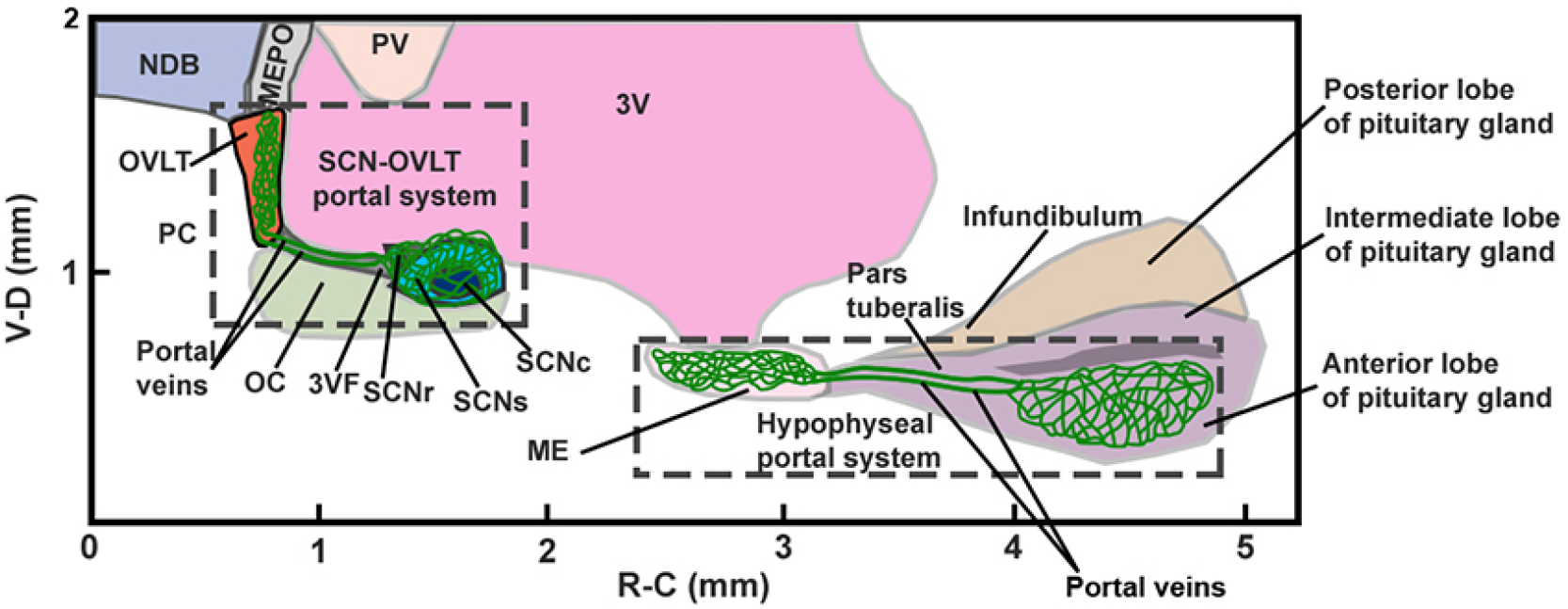
Relation of OVLT-SCN portal system to the hypophyseal portal system. The schema shows the sagittal view of the SCN-OVLT portal system and the hypophyseal portal system. Abbreviations: MEPO=median preoptic nucleus; NDB=nucleus of diagonal band; PV=periventricular hypothalamic nucleus; PC=preoptic cistern; remaining abbreviations as in Fig. 1, 2.

The sagittal view (Fig. 4) emphasizes the close proximity of the SCN and OVLT, important for effective communication of diffusible signals produced by the very small SCN nucleus (Schibler et al., 2003). Capillaries branching extensively in the AVP-rich shell region emerge from the bridge of the SCNr, and traverse the 3VF to reach the very dense capillary plexus of the OVLT (Prager-Khoutorsky & Bourque, 2015). Nearby, hypothalamic neurosecretory neurons travel to the ME, another CVO. This well-known hypophyseal portal system joins the capillary bed of the ME to the anterior lobe of pituitary gland via the pars tuberalis.

### Summary and implications

One enigma in circadian biology is how the SCN, a tiny nucleus of ~20,000 small neurons can generate a concentration of signaling proteins in body fluids that is sufficient to activate their cognate receptors. The present work points to a local and specialized target of SCN humoral signals that reach the nearby OVLT, lying at the anterior-ventral border of the 3V (Prager-Khoutorsky & Bourque, 2015). The findings establish a roadmap for exploring signaling pathways between the brain’s clock SCN and the OVLT - a sensory CVO that features a fenestrated vasculature, enriched receptors for hormones and neuropeptides and has a substantial role in the timing of drinking behavior and other physiological and behavioral functions (Gizowski, Zaelzer, & Bourque, 2016; Prager-Khoutorsky & Bourque, 2015). Likewise, these findings highlight the possibility that other CVOs bear portal systems and participate in local communications between brain, blood and cerebrospinal fluid. Late nineteenth century metaphors of the nervous system envisioned it as a syncitium on the one hand (Golgi) and as discrete neurons on the other. While Ramon y Cajal won that battle, the work in this paper however, pushes us to consider that both images are apt and that the vascular network of the brain is a key to understanding communication systems of the brain. While the road from discovery of the hypophyseal portal system to the discovery of the secretions it carried took four decades, modern tools should shorten that timeline.

## Materials and Methods

### Animals

Male C57BL/6NJ mice (The Jackson Laboratory, Bar Harbor, ME) bought at aged 8 weeks, were adapted to the lab for at least 2 weeks prior to the start of the study. Mice were provided with ad libitum access to food and water, and maintained in a 12:12-h light:dark (LD). Five hours after lights on, animals were deeply anesthetized with ketamine (100mg/kg) and xylazine (10mg/kg) and perfused intracardially with 50 ml 0.9% saline followed by 100 ml 4% paraformaldehyde (PFA) in 0.1 M phosphate buffer (PB), pH 7.3. After post-fixing in 4% PFA overnight, brains harvested for whole brain iDISCO clearing were transferred to PBS and those for brain sections immunohistochemistry were cryoprotected in 20% sucrose in phosphate buffered saline (PBS) for at least 48 hours. All procedures were carried out in accordance with the guidelines of Columbia University's Institutional Animal Care and Use Committee.

### Immunohistochemistry

Brains were sectioned (50μm) on a cryostat (Microm HM 500M, Walldorf, Germany). Free floating sections were washed in 0.1 M phosphate buffer with 0.9% saline (PBS) with 0.1% Triton X-100 (0.1% PBST), incubated in 1:100 normal donkey serum in 0.3% PBST for 1 hour, and then incubated in anti-AVP (1:5000, rabbit, Immunostar, 20069, Hudson, WI) for 1 hour at room temperature (RT) and 36 hours at 4°C on a shaker. Sections were washed in 0.1% PBST and then incubated in secondary antibody donkey anti-rabbit Cy3 (1:200, Jackson ImmunoResearch, 711-165-152, West Grove, PA) and tomato lectin-fluorescein (1:100, Vector laboratories, FL-1171, Burlingame, CA) for 2.5 hours at RT and 1 hour at 4°C on a shaker. Sections were washed with PB and mounted in PBS on subbed slides and coverslipped with Fluoromount Aqueous Mounting Medium (Sigma-Aldrich, F4680, St. Louis, MO) and cover glass No. 1 (Fisher Scientific, 12-544-18, Waltham, MA).

### iDISCO

The perfused brains were trimmed to include the olfactory tubercle to the medulla in the rostro-caudal plane and 3mm from midline in the medio-lateral plane. The iDISCO protocol of Renier et al. (2014) was used for the clearing and immunostaining with the following modifications: the tissue was incubated in the primary antibodies for two weeks and then washed for one day and incubated in the secondary antibodies for one week followed by another wash for one day and dehydration. For immunostaining the following primary and secondary antibodies were used: anti-AVP (1:5000, rabbit, ImmunoStar, 20069, Hudson, WI) with secondary donkey anti-rabbit Cy2 (1:200, Jackson ImmunoResearch,711-225-152, West Grove, PA); anti-type IV collagen (1:125, goat, SouthernBiotech, 1340-01, Birmingham, AL) with secondary donkey anti-goat Cy5 (1:200, Jackson ImmunoResearch,705-175-147, West Grove, PA); Anti-SMA (1:67, human, Dako, M0851, Santa Clara, CA) with secondary donkey anti-mouse Cy3 (1:200, Jackson ImmunoResearch, 715-165-151, West Grove, PA). AVP was used to identify the SCN. Collagen stained the whole vasculature. SMA stained for the arteries and arterioles (Kirst et al., 2020).

### Confocal microscopy

Images of horizontal and sagittal SCN vasculature were captured on a Nikon Eclipse Ti2E microscope (Nikon Inc., Melville, NY) with a 20x objective and acquired with NIS Advanced Research software. The Z stack of the horizontal image covered 23.44μm range with a step size of 0.97μm. The Z stack of coronal images covered a range of 17.18μm with a step size of 0.97μm. The Z stack of horizontal images covered a range of 24.86μm with a step size of 0.6μm. The max intensity projections are presented.

### Light sheet microscopy

Cleared, immunostained tissue was imaged using the Ultramicroscope II (LaVision BioTec, Bielefeld, Germany) equipped with a LaVision BioTec Laser Module and an Andor Neo sCMOS camera with a pixel size of 6.5 μm. The brain was mounted to be imaged horizontally with the light sheet parallel to the left-right axis. Low power images were taken first via a 0.1 NA/9.0mm WD LaVision LVMF-Fluor multi-immersion objective with a 5μm step size. The scanned volume covered the volume from the olfactory tubercle to the medulla with ~3mm from midline in both hemispheres, generating a volume of 6400μm (left-right) x 2500μm (ventral-dorsal) x 9700μm (rostral-caudal). Higher power images were taken with a 12x 0.53 NA/9.0mm WD LaVision PLAN xDISCO objective with a 2μm step size, and included the region from the OVLT to the retrochiasmatic area generating a volume of 730μm (left-right) x 1000μm (ventro-dorsal) x 1120μm (rostro-caudal). For AVP-Cy2 excitation, a 488nm, for SMA-Cy3, 525/50nm and for Collagen-Cy5 a 639nm diode laser was used. The filter sets used for detecting light emission are: 525/50nm for AVP-Cy2; 605/52nm for SMA-Cy3; 705/72nm for Collagen-Cy5.

### Image processing

The image tiles of cleared tissue were imported into Imaris (Bitplane AG, Zurich, Switzerland). The masking tool in the “Surface” module of Imaris was used to reveal the spatial relationships among SCN, OVLT, 3V and OC (Fig. 2a and Extended Data Fig. 2a), to allow simultaneous visualization of regions bearing either high and low expression levels of proteins (Extended data Fig. 1) and to distinguishSCN core and shell (Extended Data Fig. 3b).

### Vascular tracing and optic slicing

Using masked SCNc (red) and SCNs (green), the collagen labelled vasculature (Fig. 3bi) was exported to Vesselucida 360 (MicroBrightField, Colchester, VT) for tracing. The raw images for 6 SCN(s) were traced in partial projection, using the semi-manual user-guided tracing with a directional kernel method for optimal sensitivity and accuracy in detecting blood vessels. In the optic slices, step size was 50μm in the near-horizontal and 100μm in coronal slicing. The tracings of core and shell vasculature (Fig. 3bii and Extended Data Fig. 3c) were exported to Vesselucida Explorer ((MicroBrightField, Colchester, VT) for branching analysis.

### Statistical analysis

To calculate branching points in the capillary network, segments of diameter greater than 10μm were excluded (Hill et al., 2015). Data were presented as mean ± standard deviation. The volume of SCN shell is 1.53 × 10^7^ ± 1.39 × 10^6^ μm^3^ and the volume of SCN core is 3.66 × 10^6^ ± 9.16 × 10^4^ μm^3^. The normality of data was confirmed by Shapiro-Wilk test on the difference of SCN shell and core branching density (*p*=0.91) and the two-sided paired t test was conducted in R (R Core Team, 2017) to compare the density of branching points of capillaries in the shell and core of SCN, and 0.05 α level was considered statistically significant. The figure was produced with ggplot2 (Hadley Wickham, 2009). For the box plot (Fig. 3biii), the center line refers to median; the upper box limit refers to 75^th^ quantile and the lower limit refers to 25^th^ quartile; whiskers represent 1.5x interquartile range; points beyond whiskers are outliers. Dots representing measurements of the core and shell branching points from the same SCN are connected by a single line.

## Abbreviations

Abbreviations of brain regions are same as *Allen Mouse Brain Atlas* (Allen Institute for Brain Science, 2004; Lein et al., 2007) or *The Mouse Brain in Stereotaxic Coordinates* (Paxinos & Franklin, 2012).

## Reporting summary

Further information on research design is available in the Nature Research Reporting Summary linked to this paper.

## Data availability

Datasets supporting the findings of this paper are available from the corresponding authors upon reasonable request.

## Acknowledgements

We thank K.L. Olsen and A.J. Silverman for comments on earlier drafts of the manuscript; C. McKernan for technical support; R. Tomer for advice on image processing, P. Buchanan for assistance with animal care. Light sheet microscopy was performed with support from the Zuckerman Institute’s Cellular Imaging platform, and the National Institute of Health (NIH) grant 1S10OD023587-01. We are grateful for support of this work by National Science Foundation (NSF) grant 1749500 (to RS).

## Author contributions

Y.Y., J. L. and R.S. designed the experiment. Y.Y. and R.S. wrote the manuscript. J.L. and A.B.T. commented on early drafts of the manuscript. A.B.T. and Y.Y. acquired light sheet microscope images and prepared figures. A.B.T. conducted confocal microscopy and light microscopy. Y.Y. conducted tissue clearing, image processing, tracing and statistical analysis.

## Additional information Supplementary information

TBD

## Correspondence and requests for materials

should be addressed to Y.Y or R.S.

## Peer review information

TBD

