## Supplemental material for "The brain clock portal system: SCN-OVLT"

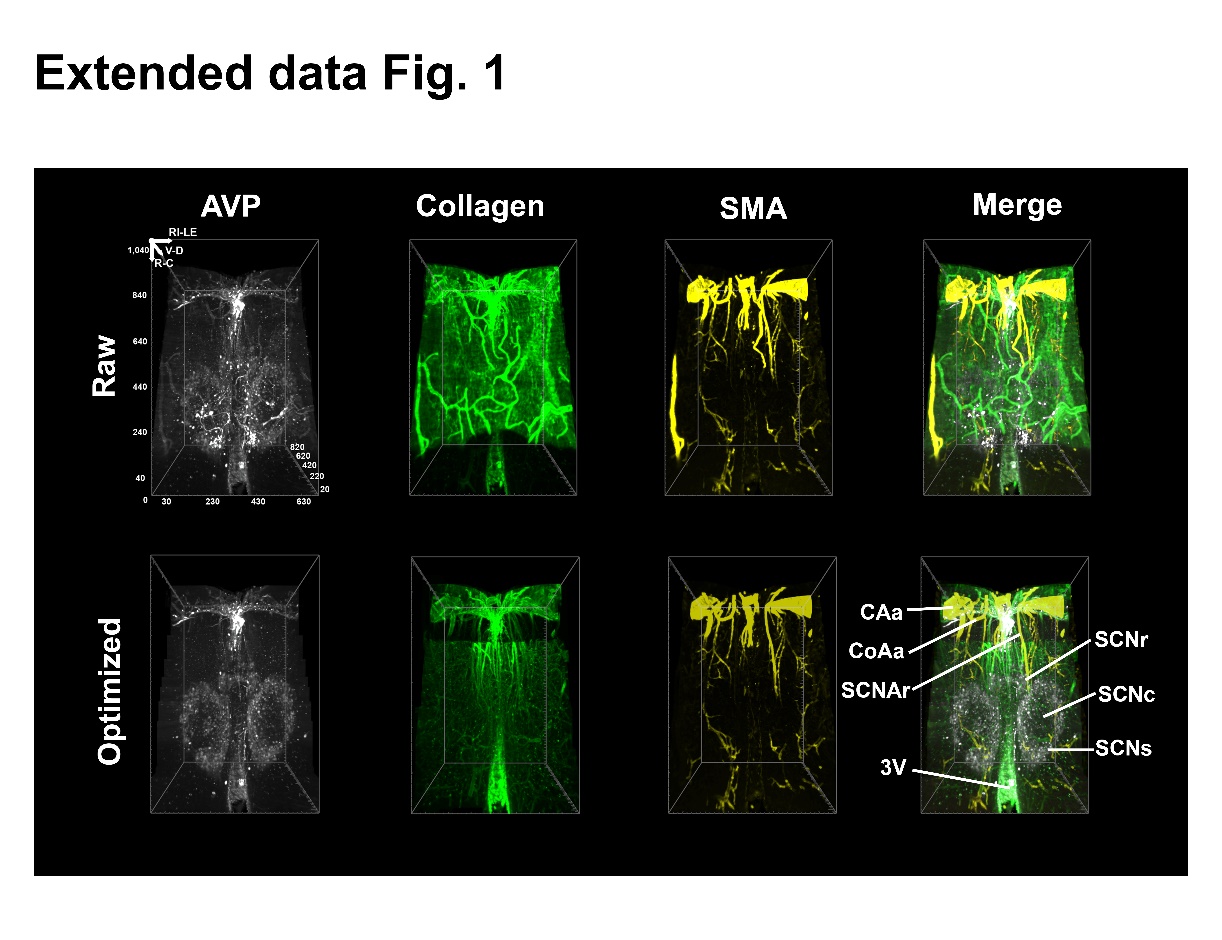

**Extended Data Fig. 1 | Blood vessels communicating between SCN-OVLT.** The raw images of AVP, collagen and SMA are shown in the upper row. In the optimized channels, masking was used for better visualization of very intense and less intense label in the same image (e.g. collagen in OVLT vs SCN) and to eliminate signals from the pia and OC. AVP (white) demarcates the SCN main body and the rostrum which lies in the midline and forms the rostral pole of the nucleus. Reference axis: C=caudal; R=right; D=dorsal. Scale unit=µm. In the collagen channel (green), a rich bundle of blood vessels lies along the midline. SMA (yellow) labels the arteries in the SCN area. The merged image shows that the midline blood vessels travel between the OVLT and SCN. Arteries branching from the ACA and ACoA which supply the SCN are lateral to the midline blood vessels. Voxel=0.54µm (rostro-caudal) x 0.54µm (left-right) x 2µm (dorso-ventral). Abbreviations: SCNAr=SCN artery, rostral branch; remaining abbreviations as in Fig. 1, 2.

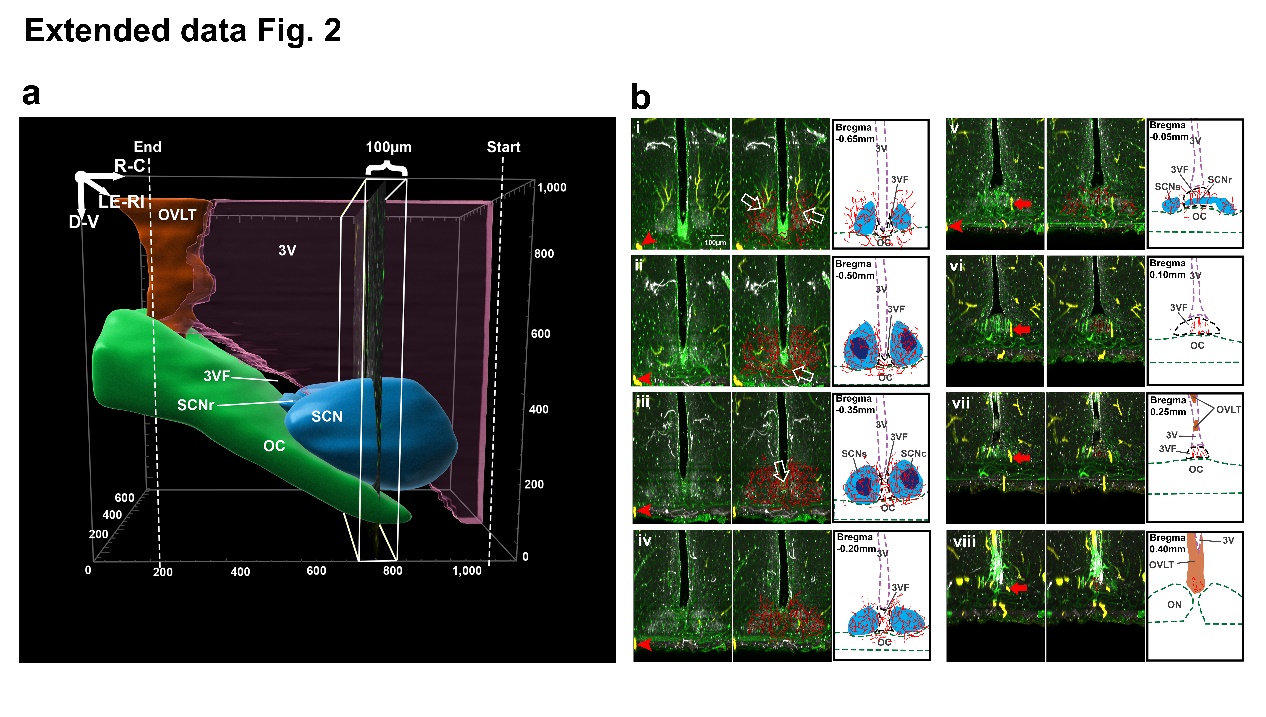

**Extended Data Fig. 2 | Capillaries connecting OVLT and SCN in coronal optical scans. a,** For analysis of the vasculature of areas of interest, masking was used as in Fig. 2a. The white rectangle shows the orientation of the coronal serial scans made from the SCN to the OVLT starting from the caudal aspect of the SCN. Reference axis: C=caudal; L=left; V=ventral. Scale unit=µm. **b,** Vasculature in serial coronal optic slices from Bregma -0.65 to 0.40. The plates show triplets of images as follows: left panel= merged AVP, collagen and SMA; red arrows and arrowheads are the place markers indicating landmarks for orientation in adjacent slices. Middle panel=blood vessel traces are superimposed on immunochemical results of the left panel. Right panel=drawing identifying structures in the middle panel. Color coding as in Fig. 2c. Detailed descriptions of image sequences as follows: **bi,** In the caudal-most SCN, arteries are running vertically through the SCN (middle column, white arrows). **bii-iv**, The SCN vasculatures between hemispheres are connected by capillaries in the chiasm (bii**,** middle column, white arrow) and in the 3VF (biii, middle column, white arrow). **bv,** Blood vessels from bilateral SCNs form anastomoses near SCNr. **bvi-vii,** Portal blood vessels travel in the 3VF. **bviii,** Portal blood vessels join OVLT capillary network from its bottom. Slice thickness=100µm. Abbreviations as in Fig.1, 2.

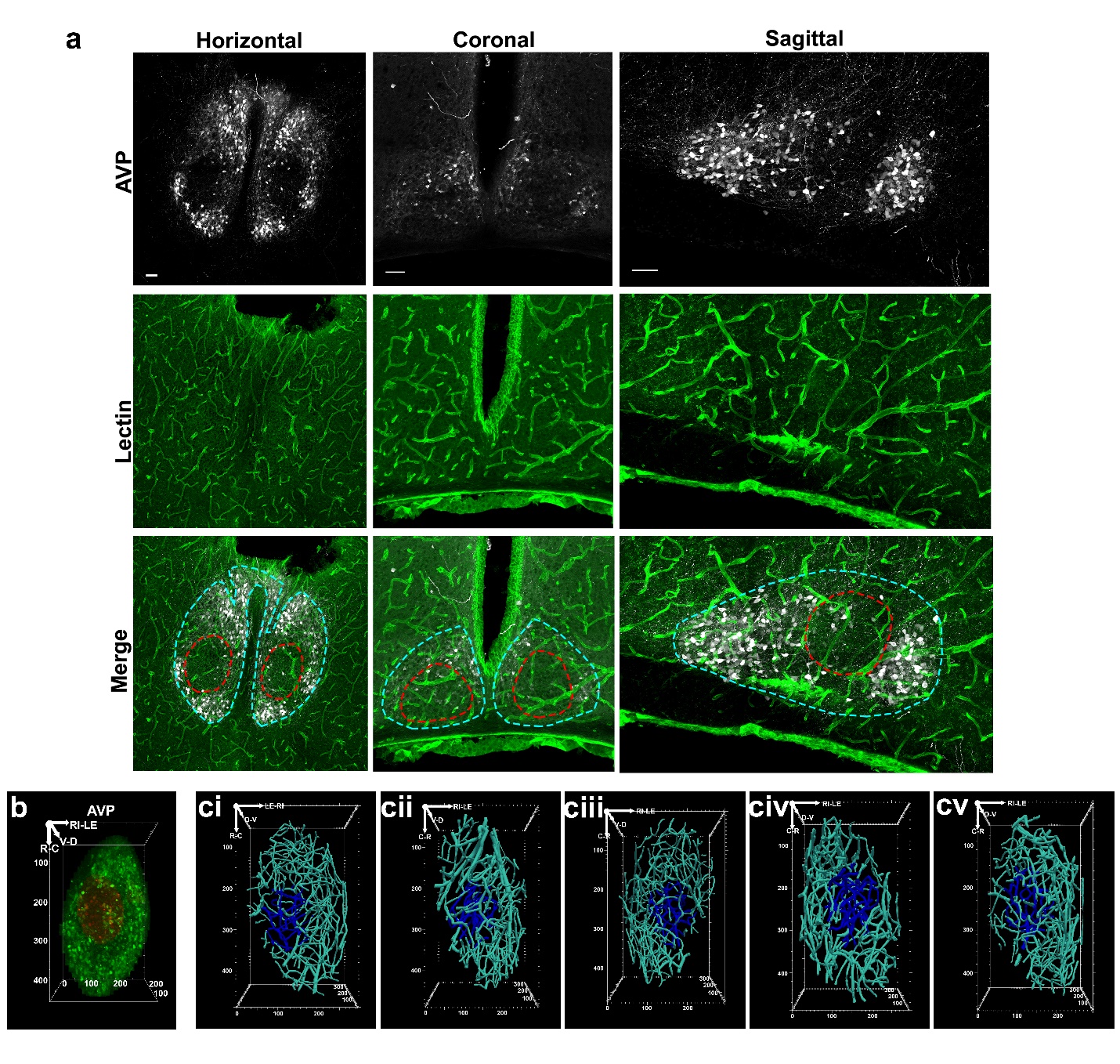

**Extended Data Fig. 3 | SCN shell vs. core. a,** Confocal microscopic images of SCN in three orientations distinguishes the AVP poor core from the AVP rich shell. Merged images, in each orientation, suggest that there is greater vasculature density in the SCN shell (dashed blue line) compared to the core (dashed red line). Best seen in the AVP-labelled horizontal view is the SCNr which lies right at the midline, the locus of capillary blood vessels connecting SCN and OVLT. Scale bar=50µm. **b,** AVP staining after masking shows shell (green) and core (red) in a representative SCN prepared for vasculature extraction. Reference axis: R=rostral; L=left; D=dorsal. Scale unit=µm. **c,** The statistical analysis in Fig. 3biii was based on tracings of SCN(s) in Fig. 3bii and ci-v. Color coding as in Fig. 3b. Reference axes: R=rostral, L=left, and D=dorsal in ci; ciii-v. C=caudal, R=right, and D=dorsal in cii. Scale unit=µm.
